# Low nucleus to cytoplasmic ratio compromises cytokinesis, genome stability and development in haploid embryonic cells

**DOI:** 10.1101/2025.08.30.673237

**Authors:** Michael V. Zuccaro, Daniela Georgieva, Shuangyi Xu, Matthew Dennen, Qian Du, Gloryn Chia, Selma Amrane, Lisa C. Grossman, Alejandro De Los Angeles, Robert Prosser, Rogerio Lobo, Dieter Egli

## Abstract

Mammalian species exist as diploid organisms, with only gametes existing as stable, nonproliferating haploids. Haploid mammalian pluripotent stem cells, in all derived species to date - mouse, human and rat - undergo spontaneous diploidization. Here, we investigated the mechanisms affecting the stability of the haploid state using the mouse and human embryo, as well as a human embryonic stem cells. We demonstrate that diploidization occurs early in preimplantation development and is often unproductive, with haploid embryos exhibiting decreased developmental potential. Haploid embryos show increased chromosome segregation errors at the first mitosis, failure to align chromosomes on the metaphase plate in the second mitosis. Delayed mitotic progression and failure to form a central spindle and a midbody are followed by cytokinesis failure, diploidization, increased DNA damage marked by γH2AX and RPA32 foci, and low developmental potential. By increasing the nucleo-cytoplasmic ratio in diploids, or reducing this ratio in haploids, these phenotypes can be induced in diploids or reduced in haploids. Changing ploidy alone from haploid to diploid without also adjusting nucleo-cytoplasmic ratio does not improve mitotic phenotypes and developmental outcomes. Thus, the most upstream driver responsible for the instability of the haploid state is a stage-specific ratio of nucleus to cell size.

## INTRODUCTION

Haploid embryonic stem cells have been derived from multiple mammalian species including mouse (Leeb and Wutz, 2011, Yang et al., 2012, Elling et al., 2011), rat (Li et al., 2014), and human (Sagi et al., 2016). Haploid derivation via parthenogenesis in mammals is achieved through the activation of an unfertilized oocyte without sperm, whereas androgenesis is achieved by injection of a sperm into an oocyte and removal of the maternal genome. In both instances of uniparental derivation, haploid embryonic development proceeds to the blastocyst stage, enabling the derivation of embryonic stem cells, albeit at reduced efficiency compared to fertilized diploids, and only a portion of the resulting lines contain haploid cells (Sagi et al., 2016, Sagi et al., 2019, Ding et al., 2015). The haploid cells can be purified through sorting and essentially pure haploid cell lines can be isolated (Sagi et al., 2016). Haploid lines continue to undergo diploidization at a rate of ∼3% per population doubling at low passage, acquiring a second set of all chromosomes, including the X chromosome. Unless routinely selecting for haploids via cell sorting by DNA content (Leeb and Wutz, 2011) or cell straining for size (Qu et al., 2018, Freimann and Wutz, 2017), haploid cells will gradually and irreversibly convert to a diploid state, homozygous for all chromosomes. The low stability of haploid cell cultures and conversion to diploids is a well-documented phenomenon in all species examined, including in mouse (Kaufman et al., 1983, Leeb and Wutz, 2011), rat (Li et al., 2014, Hirabayashi et al., 2017), and human (Sagi et al., 2016, Zhang et al., 2020). Using live cell microscopy, mitosis via failed cytokinesis or mitotic slippage is thought to be the primary cause of diploidization in human embryos (Leng et al., 2017).

Changes in ploidy are not unique to haploid pluripotent stem cells, as polyploidy is observed in various cell types and organs, including brain cells, liver cells, placenta, and macrophages, as reviewed in (Sagi and Benvenisty, 2017). One of the most studied organs with regards to ploidy is the liver, in which a failed cytokinesis event yields binucleate tetraploid cells. These cells convert to tetraploid mononucleate hepatocytes at the following mitosis (Guidotti et al., 2003). Failure of cytokinesis involves the lack of the formation of a midbody, even as two groups of chromosomes do separate from each other (Celton-Morizur et al., 2009). Tetraploidization is mediated by the AKT-PI3 kinase pathway downstream of insulin signaling. Reduction of insulin levels or inhibition of PI3K results in reduced cytokinesis failure and fewer binucleated cells.

Polyploidy in cardiomyocytes also progresses through mitosis without cytokinesis followed by binucleation (Leone et al., 2018). During tumorigenesis, increases in ploidy are common and may facilitate the tolerance to new mutations and the development of cancer (reviewed in (Davoli and de Lange, 2011)). Thus, changes in ploidy have broad relevance to developmental processes as well as disease. Haploid embryos and stem cells provide a model to study ploidy change, with the most direct relevance to developmental processes in the placenta where ploidy changes also occur.

Here we show that, in both mouse and human systems, diploidization occurs early in development and is observed with an increase in binucleated cells, delayed mitotic entry and progression, and failed cytokinesis. Failure of cytokinesis is associated with the lack of metaphase alignment, failure to form a central spindle and failure of Aurora B and Survivin localization to the midbody. In developing embryos, we find that most diploidization events are unproductive and due to cell cycle delay and arrest. DNA damage markers _γ_H2AX and RPA32, indicating double strand breakages and single-stranded DNA regions, are increased in haploid mouse embryos at the 4-cell stage. Analysis of binucleated cells in diploidizing embryos reveals a high degree of aneuploidy, primarily due to whole chromosome segregation errors. CHK1 is upregulated in haploid human ESCs, and its inhibition alleviates mitotic delays in both haploid human ESCs and haploid mouse embryos. Through a series of embryonic manipulations, it was determined that failure of cytokinesis is driven in haploids by a low nuclear-cytoplasmic ratio, rather than by ploidy alone. Increasing this ratio in haploid mouse embryos improves mitotic timing and progression, and developmental outcomes, whereas decreasing the nuclear-cytoplasmic ratio in diploid control embryos induced mitotic delays, cytokinesis failure and poor developmental outcomes. We thus show that the ratio of nucleus to cytoplasmic content ratio is the primary determinant of genome instability in haploid mammalian cells, contributing to diploidization.

## RESULTS

### Diploidization in human embryos at the cleavage stage includes binucleation and delayed diploidization

Haploid mammalian embryonic stem cells are challenging to derive due to the conversion to diploid cells in the embryo prior to derivation, and after derivation in culture. Conversion to diploidy is unidirectional – there is no spontaneous return to haploidy. Combined with the relative stability of the diploid state, the conversion from haploid to diploid is gradual, and eventually complete. Thereby these cells provide a model to study ploidy changes and the mechanisms determining stable ploidy.

To determine at which point during human haploid development diploidization occurs, human parthenogenetic embryos were artificially activated by an electrical pulse. Human androgenetic embryos were activated via enucleation of the oocyte’s nucleus and injection of a sperm cell.

Both types of uniparental embryos were analyzed for ploidy at various developmental stages, until the blastocyst stage. Ploidy was assessed via immunofluorescent staining of the centromere; each chromosome will have a single centromere and as such, the number of centromere foci can be used for determining ploidy in embryonic stem cells (**Fig. 1A**) and human embryos (**Fig.1B**) (Sagi et al., 2016). 23 chromosomes, and thus centromeres, are expected in haploid nuclei and 46 in diploids. Parthenogenetic and androgenetic embryos exhibited binucleation at every stage of development toward the blastocyst, suggesting a failure to complete cytokinesis and produce separate daughter cells, each with a with single nucleus (**Fig. 1C, D**). Despite the presence of binucleate cells as early as the cleavage stage and throughout subsequent developmental stages, the presence of mononucleate diploid cells were not observed until the morula and blastocyst stage on day 4 and after of development (**Fig. 1C, D**). The proportion of mononucleate diploids in subsequent developmental stages relative to when binucleate cells are first observed indicates that most binucleate cells do not convert to mononucleate diploids, and that these diploidization events are unproductive mitotic events without contribution toward further development.

**Figure 1.**
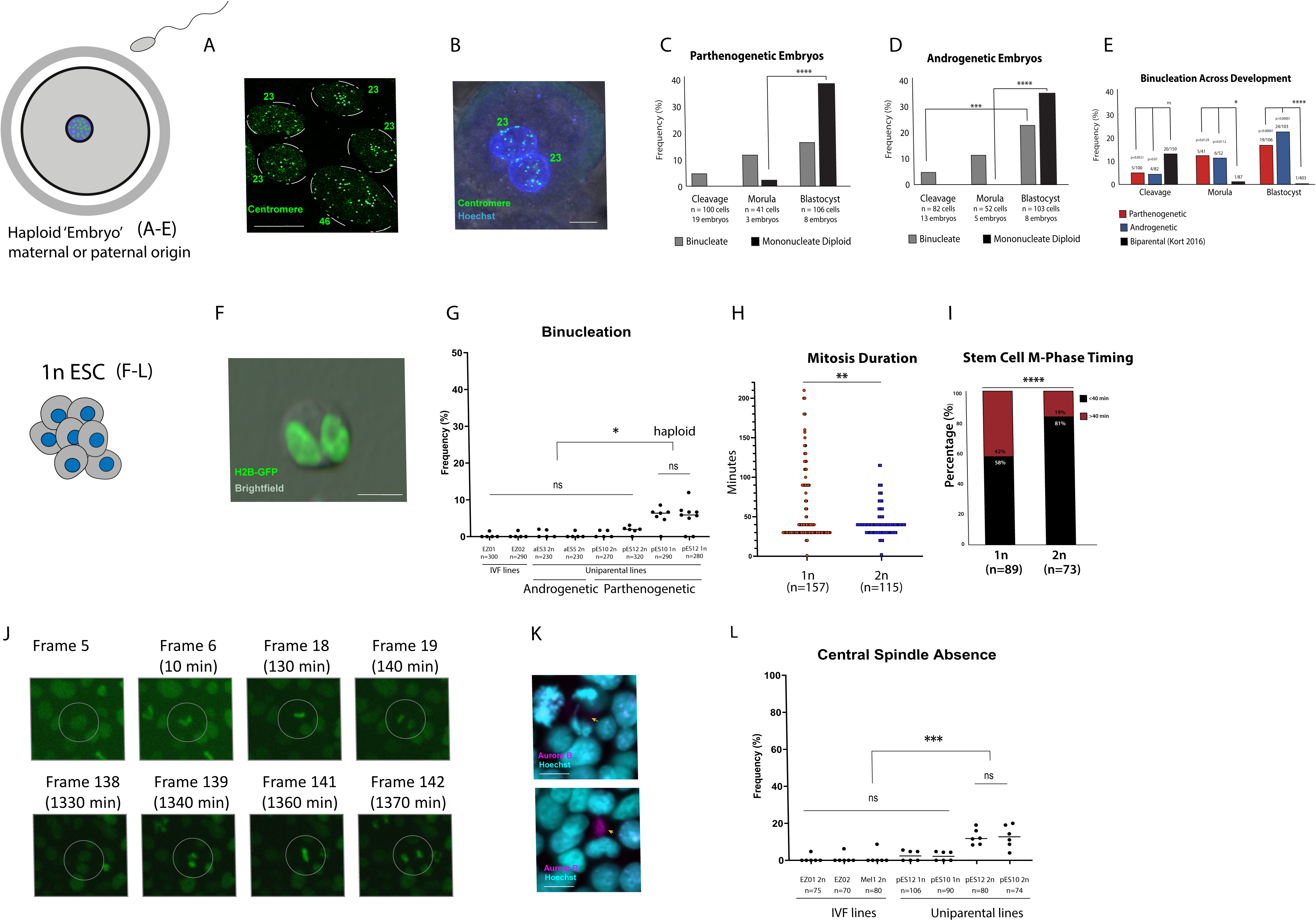
Haploid human embryos exhibit binucleation and unproductive diploidization events as early as the cleavage stage. (**A**) Immunostaining of centromeres in nuclei of androgenetic human embryonic stem cells. Nuclei edge labeled with dashed line, centromere counts indicated (haploid = 23; diploid = 46). (**B-D**) 69 embryos were assessed via confocal microscopy, including 36 parthenogenetic (8 2-4 cell, 11 cleavage stage, 6 multi-cell, 3 morula, 8 blastocyst TE biopsy) and 33 androgenetic (5 2-4 cell, 8 cleavage stage, 7 multi-cell, 5 morula, 8 blastocyst TE biopsy). (**B**) Immunostaining of a day 3 embryo for centromeres. Note the binucleate cell, each nucleus with 23 centromeres. Scale bar = 10 μm. (**C**) Quantification of the frequency of binucleate vs mononucleate diploid cells determined by immunostaining for centromeres, in parthenogenetically derived human embryos at multiple developmental stages. (**D**) Quantification of frequency of binucleate vs. mononucleate diploid cells determined by immunostaining for centromeres, in androgenetically derived human embryos at multiple developmental stages. (**E**) Binucleation across developmental stages for parthenogenetic, andrognetic, and IVF-derived embryos. (**F**) Representative image of binucleation in a haploid human embryonic stem cell transfected with H2B-GFP. Scale bar = 10 μm. (**G**) Quantification of binucleation from fluorescent images of haploid and diploid human embryonic stem cells. Scale bar = 10 μm. One-way ANOVA; *=p<0.05. Data taken from 6 independent experiments. (**H**) Mitotic duration in haploid and diploid ESCs. (**I**) Quantification of cells with mitotic timing of greater than 40min. (**J**) Time lapse microscopy of H2B-GFP labeled ESCs showing delay. (**K**) Representative immunofluorescent imaging of the midbody (Aurora B), and DNA (Hoechst 33342) in human embryonic stem cells. Scale bar = 10 μm. (**L**) Quantification of central spindle formation in haploid and diploid human embryonic stem cells evaluated by Aurora B immunostaining ana-/telo-phase cells. Data from 3 independent experiments. Fisher exact test; **=p<0.01.

Haploid embryos, both parthenogenetic and androgenetic exhibited significantly higher frequencies of binucleation at the morula and blastocyst stages compared with biparental embryos, but without significant difference in frequency between the parent of origin (parthenogenetic vs androgenetic) (**Fig. 1E**). These findings of increased binucleation are consistent with earlier studies of haploid parthenogenic human embryos (Leng et al., 2017), while our studies also include androgenetic embryos, which contain a sperm-derived centrosome. Thus, binucleation in parthenotes is not due to the absence of a centrosome, but the consequence of haploidy.

### Haploid human embryonic stem cell populations exhibit subsets of delayed M phase and decreased midbody formation

To determine whether diploidization in ESCs also involves binucleation, haploid ESCs and isogenic diploidized cell lines were stably transfected with H2B-GFP, in addition to two newly derived diploid biparental control ESC lines (**Fig. S1**). Haploid human stem cell lines exhibited an increase in binucleate cells compared to isogenic diploidized cells, and in biparental diploids (**Fig. 1F, G**). In addition to parthenogenetic lines, androgenetic embryonic stem cell lines were also included to determine whether these results are due to the haploid state or due to parental origin (maternal vs paternal). There was no statistically significant difference between parthenogenetic and androgenetic diploid cell lines and biparental diploid cell lines (**Fig. 1G**). suggesting that the rate of binucleation and thus diploidization are a consequence of the haploid state rather than due to uniparental contribution.

Haploid and isogenic diploid ESCs transfected with H2B-GFP were used for live cell imaging. Observing the duration between chromosome condensation and daughter nuclei separation revealed significant differences between haploid and diploid M phases. Analysis at 10-minute intervals showed that diploids typically exhibited a mitosis within 40 min (**Fig. 1H**), with less than 20% of cells taking slightly longer to complete mitosis (**Fig. 1I**). While haploids exhibited many cells completing mitosis within 40 minutes, 58% of haploids required more time (**Fig. 1I**). Delays in haploids ranged from 50 minutes to upwards of 3 hours to complete M phase (**Fig. 1H**). These data show that a subset of haploid human embryonic stem cells exhibit a prolonged M phase. A prolonged M phase was also observed in parthenogenetic human embryos (Leng et al., 2017) and mouse embryonic stem cells (Guo et al., 2017). A prolonged M phase can result in failed cytokinesis, a binucleate diploid cell, and at the next mitosis, a conversion to a mononucleate diploid cells (**Fig. 1J**).

Haploid human embryonic stem cells were analyzed for abnormalities in mitosis. Staining of two genetically distinct haploid stem cell lines (pES10 and pES12) revealed that both lines exhibited a decrease in midbody formation (89% and 86%, respectively) compared to isogenic diploid lines (97-100%), as well as biparental, IVF-derived lines (100%) (**Fig. 1K**, **L**). M phase delays and lack of a midbody followed by failed cytokinesis and by binucleation contributes to diploidization.

### Haploid mouse embryos exhibit chromosome segregation errors and abnormal mitotic progression

To study diploidization at a single cell level in more detail, we used the mouse embryos. The mouse embryo has the advantage that it begins from a normal haploid state, and the the chromosome content can be manipulated. Furthermore, the well-documented timing of the cell cycle across mouse embryonic development allows for precise and accurate monitoring of mouse embryos and mitotic outcomes during these first two cell cycles. Mouse oocytes were activated and haploid and diploid embryos generated through conditions permitting or suppressing polar body extrusion (**Fig. 2A**), resulting in interphase 1-cell embryos with either one or two maternal pronuclei (**Fig. 2B**). Spindle assembly and chromosome segregation was observed at the first mitosis 14-16h post activation and at the second mitoses (32-40h post activation) via brightfield microscopy and live-cell imaging (**Fig. 2C**). Mitotic entry was identified by the moment of pronuclear envelope breakdown (PNEBD). PNEBD occurred in 100% of mouse embryos at the first mitosis for both haploid and diploid groups, with identical timing between groups (**Fig. 2D**). Similarly, the time from PNEBD to cleavage during the first mitosis was identical in haploid and diploid mouse embryos (**Fig. 2D**). Haploids efficiently cleaved to two cells (n=163) except for a single haploid egg (**Fig. 2D**).

**Figure 2.**
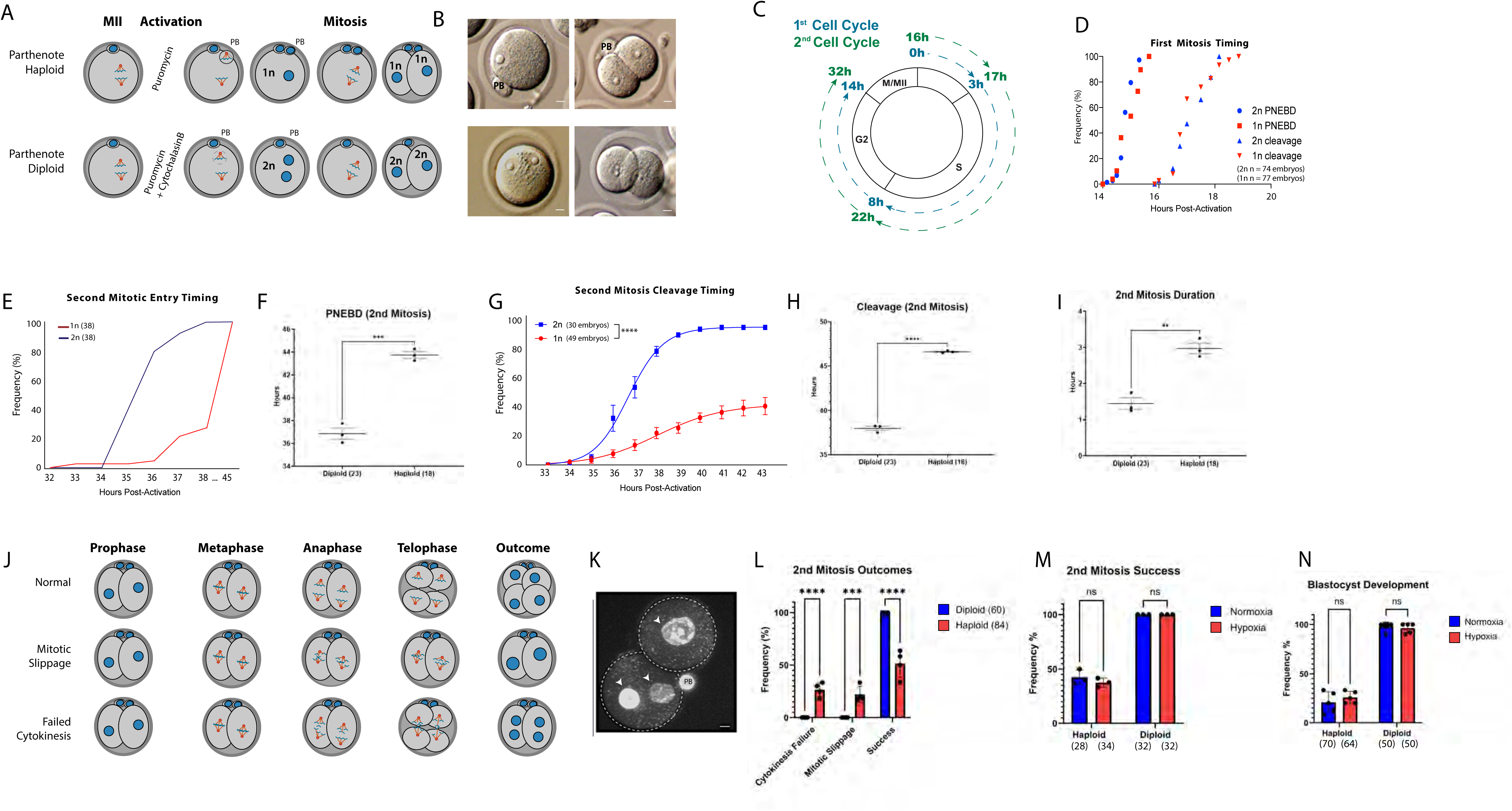
Haploid mouse embryos have poor developmental potential and frequently exit mitosis via mitotic slippage and failed cytokinesis (**A**) Illustration of mouse haploid and diploid embryo development post-activation. (**B**) Haploid (top) mouse embryos including visible polar body (labeled PB) at both 1- and 2-cell stages and diploids (bottom), including binucleation at the 1-cell stage. Scale bar = 10 μm. (**C**) Schematic illustrating the timing of the first and second mitoses in activated mouse embryos in hours post activation. (**D**) Timing (hours post activation) of mitotic entry and cleavage at the first mitosis in haploid (n=77) and diploid (n=74) mouse embryos. (**E**) Mitotic entry timing (hours post activation) of haploid and diploid embryos at the second mitosis. (**F**) Timing of pronuclear envelope breakdown based on live-cell imaging during the second mitosis taken at 10-minute intervals. T-test performed; p<0.0001. (**G**) Haploid and diploid cleavage timing (hours post activation). Fisher exact test performed at final timepoint. (**H**) Timing of second mitosis based on live-cell imaging. T-test. (**I**) Duration of 2^nd^ mitosis in haploid and diploid mouse embryos. (**J**) Schematic for a normal mitosis, mitotic slippage and failed cytokinesis at various stages of mitosis. (**K**) Image of Hoechst 33342 staining depicting failed cleavages resulting in a binucleate (two arrowheads) cell and mononucleate (single arrowhead) cell. PB= polar body. Scale bar = 10 μm. (**L**) Quantification of cleavage outcomes at the second mitosis in 60 haploid cells and 84 diploid cells from 4 independent experiments. (**M**) Quantification of cleavage success in hypoxic and standard oxygen concentrations. Fisher exact test. (**O**) Quantification of successful blastocysts development in haploid and diploids from 5 different experiments. Fisher exact test.

In stark contrast, haploids exhibited a noticeable difference in the timing of both mitotic entry and progression. The time until the pronuclear envelope breakdown during the second mitosis was delayed in haploid mouse embryos compared to diploid mouse embryos by 4-6 h (**Fig. 2E, F**). Similarly, the time from pronuclear envelope breakdown until cleavage was also delayed in haploid mouse embryos compared to diploids (**Fig. 2G, H**). Though all haploids entered mitosis, the majority of haploids failed to undergo cytokinesis (**Fig. 2G**). Haploid embryos spent on average 2 more hours in mitosis compared to diploids (**Fig. 2I**). Abnormal mitotic outcomes included binucleation and diploid mononucleation (**Fig. 2J, K, L**). In 84 cells, all of which entered mitosis, 58% of failed cleavages were a result of failed cytokinesis, and 42% were a result of mitotic slippage, distinguishable by the presence of either binucleation or a single large nucleus, respectively (**Fig. 2L**). Overall cleavage success was significantly lower in haploid embryos (51%) compared to diploid embryos (100%) at the second mitosis (**Fig. 2L)**. An earlier study suggested that oxygen tension affected diploidization (Zhang et al., 2020), but in this study, 4-5% oxygen had no effect on successful cleavage at the second mitosis (**Fig. 2M**).

Haploid mouse embryos exhibited a frequent failure of blastocyst development; most of the embryos arrested, with only 19% developing to the blastocyst stage, in contrast to diploids exhibiting over 90% blastocyst development. Haploid development to the blastocyst stage was reduced relative to diploids independent of oxygen tension (**Fig. 2N**).

### Haploid mouse embryos lack central spindle and midbody formation

To understand the molecular defects leading to cytokinesis failure, we sought to observe chromosome segregation outcomes as well as central spindle and midbody formation. There was a slight, albeit significant increase in the frequency of chromosome segregation errors at anaphase of the first mitosis of haploids compared to diploids, observable by immunofluorescent staining of chromosomes with phospho-histone H3 and Hoechst (**Fig. 3A,B)**. At the second mitosis, haploid embryos exhibited substantially more chromosomes segregation errors, marked by lagging chromosomes, when compared to diploids, and also compared with the frequency of errors observed in the first mitosis (**Fig. 3C**). Among chromosome segregation errors, chromosome fragmentation was also observed upon immunofluorescent staining (**Fig. 3C**). Poor or entirely absent metaphase alignment was also observed in haploids at the second mitosis (**Fig. 3D**). Survivin and β-tubulin was used to visualize midbody formation. In the first mitosis, haploid embryos did not significantly differ from diploids in terms of central spindle and midbody formation, as both groups exhibited normal Survivin immunostaining in all mitoses observed (**Fig. 3A**). In contrast, during the second mitosis, the rate of failure to form a midbody was significantly higher in haploids compared to diploids (**Fig. 3E**). All diploids formed midbodies, compared to only 31% in haploids (**Fig. 3F**).

**Figure 3.**
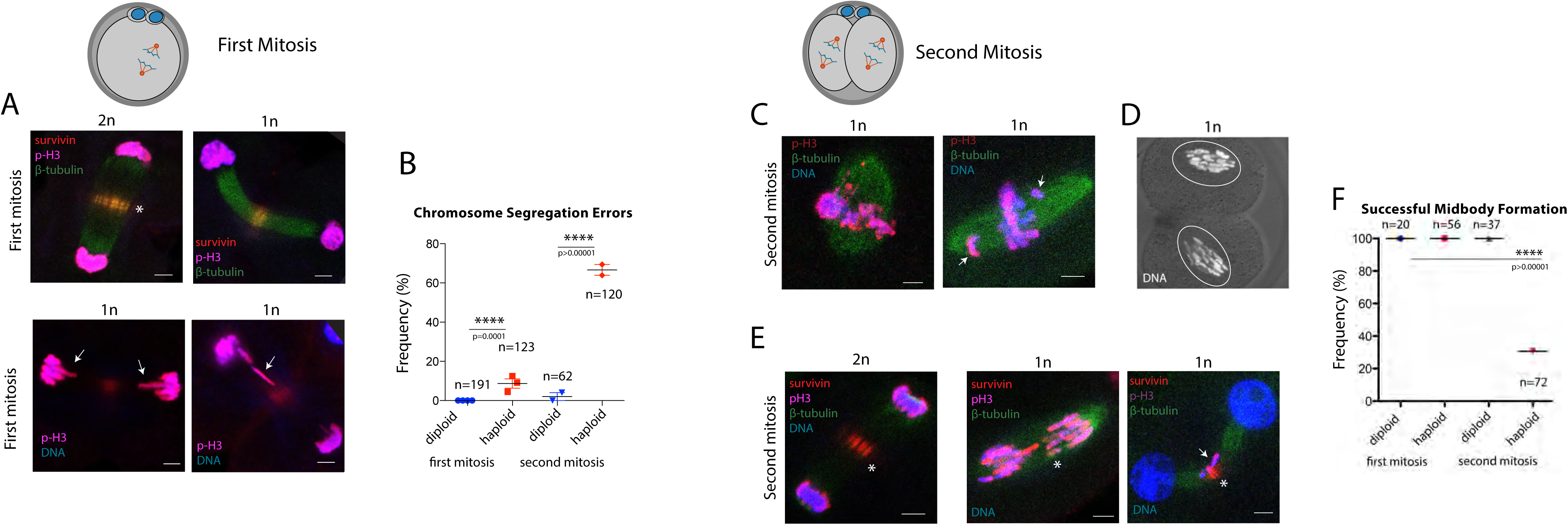
| Haploid mouse embryos fail central spindle and midbody formation (**A**) Immunofluorescent staining for pH3, Survivin, and spindles (β-tubulin), of normal mitosis of diploid (2n) and haploid (1n) mouse embryos at the first mitosis. Scale bar = 10 μm. (**B**) Quantification of chromosome segregation errors, identified by abnormal anaphase. Fisher’s Exact Test. (**C**) Haploid embryos at the second mitosis exhibiting chromosome fragmentation, poor chromosome condensation, and improperly aligned chromosomes. (**D**) Haploid mouse embryos at the second mitosis with lack of metaphase alignment despite a fully formed spindle visible by the granule-free area (outline). (**E**) Representative images of the second mitosis in a diploid and haploid mouse embryos. (**F**) Quantification of the frequency of midbody formation, identified by β-tubulin and Survivin localization in mouse embryos. Fishers Exact Test.

### Frequent whole chromosome segregation errors in mouse haploid embryos

To better understand the cause of abnormal chromosome segregation, karyotype analysis was performed at the 4-cells stage and its binucleate equivalent of half the cell number (**Fig. 4A**). In binucleate cells, individual nuclei were isolated (n=19), amplified and analyzed separately, and individual blastomeres of cleaved 4-cell embryos (n=67) were also collected (**Fig. 4A**). 104 blastomeres from 26 diploid fertilized control embryos were also evaluated (**Table S1**). Haploid nuclei from binucleate cells, as well as blastomeres from cleaved 4-cell embryos showed frequent aneuploidies involving one or several chromosomes (**Fig. 4B, C**). Whole chromosome segregation errors were most frequently observed: of a total of 89 abnormal chromosomes, only 2 were segmental, and 3 showed multiple breakages. Surprisingly, two thirds of the nuclei in cells with failed cytokinesis were euploid. Segregation errors were equally frequent in binucleate cells as in cleaved blastomeres (31 vs. 29%), and significantly different from diploid controls (**Fig. 4D**), concordant with cytological observations. Thus, cytokinesis failure can occur despite bipolar attachment and accurate chromosome segregation, and abnormal segregation does not necessarily lead to cytokinesis failure.

**Figure 4.**
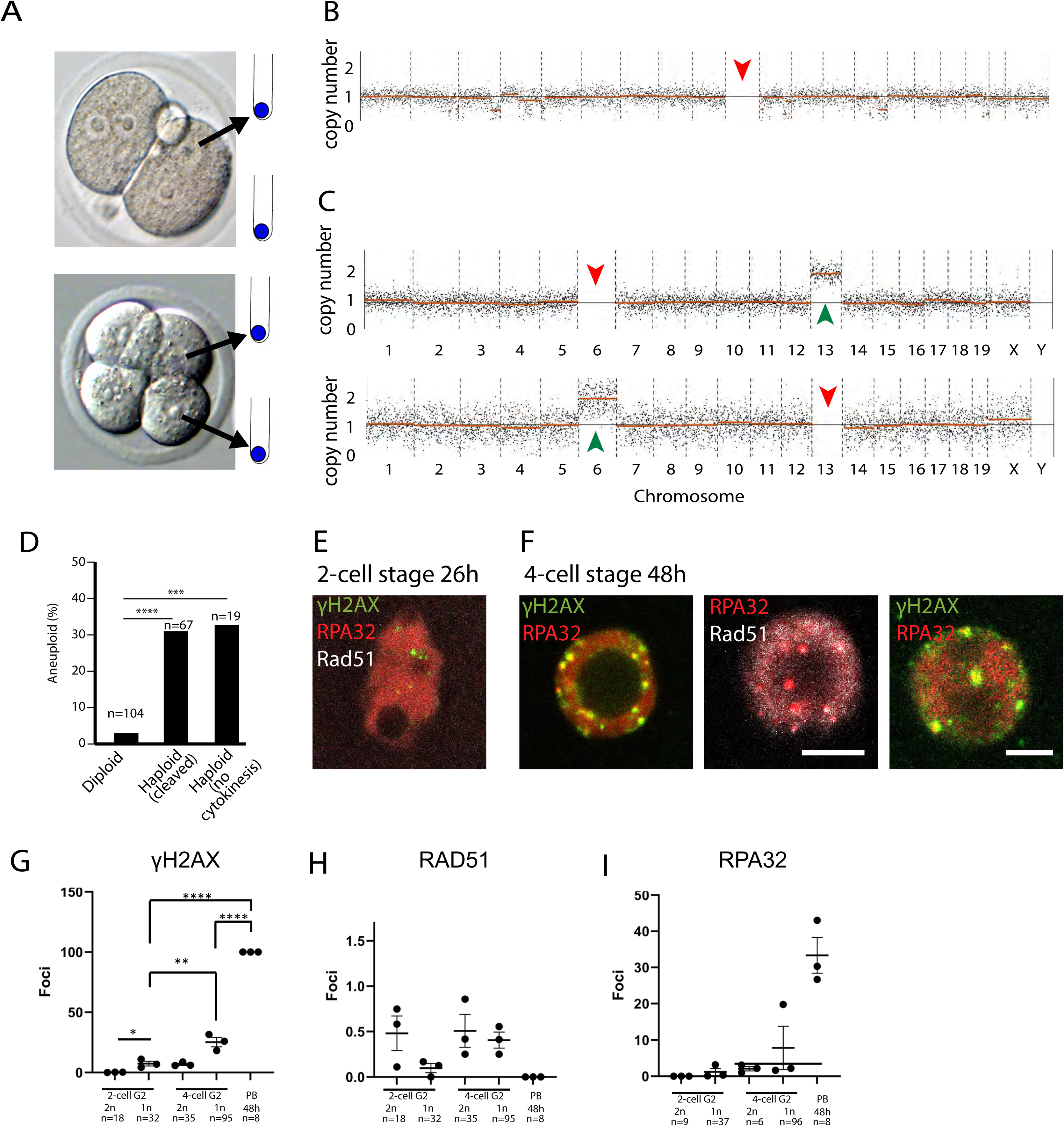
Haploid mouse embryos exhibit increased chromosome segregation errors and DNA damage (**A**) Binucleate and cleaved haploid mouse embryos after the second mitosis, equivalent to the ‘4-cell’ stage. Individual nuclei or blastomeres are collected and analyzed for chromosome content using NGS. (**B**) Copy number karyotype plot illustrating reciprocal whole chromosome exchange of a nucleus from a binucleate blastomere. Arrow indicates loss. (**C**) Karyotype plots of sister blastomeres showing complementary chromosome segregation errors. Note complementary losses and gains indicated by arrows. (**D**) Quantification of chromosomal aneuploidy in binucleate blastomeres and cleaved 4-cell haploids and controls. **** p<0.00001, *** p< 0.0001. Fisher exact test. (**E, F**) Immunostaining for indicated DNA damage markers All scale bars = 10 μm. (**E**) 2-cell haploid mouse embryo. (**F**) 4-cell stage haploid mouse embryos. (**G**) Quantification of γH2AX foci. (**H**) Quantification of RAD51. (I) Quantification of RPA32 foci.

### DNA damage foci are increased in haploid mouse embryos at the 4-cell stage in G2 phase

Segmental chromosome errors in haploid mouse embryos suggest that some chromosome missegregation events involve DNA damage. We analyzed mouse embryos at precisely defined time points, at 28h and 48h post-activation, or in G2 of the 2-cell stage, and in G2 of the 4-cell stage (or its developmental equivalent), as these timepoints are before and after when frequently observed chromosomal segregation errors would occur. Immunofluorescent staining was performed for _γ_H2AX, RPA32, and RAD51 (**Fig. 4E, F**). Staining was performed in both haploid and diploid embryos. At G2 of the 2-cell stage, _γ_H2AX foci were increased in haploid embryos compared to diploids (**Fig. 4G**), but there was no significant difference between RPA32 or RAD51 foci (**Fig. 4H, I**). The frequency of DNA damage foci _γ_H2AX foci and RPA32 increased from the 2-cell to the 4-cell stage in both haploids and diploids. There was a significant increase in _γ_H2AX and RPA32 foci, but not RAD51 foci in haploids at the 4-cell stage compared to diploids at the 4-cell stage (**Fig. 4G-I**). As a point of reference for positive staining, the polar body exhibited a large number of foci for DNA damage markers _γ_H2AX and RPA32 (**Fig. 4G, I**), but none for RAD51 (**Fig. 4H**), suggesting insufficient DNA repair in the polar body. There was a significant decrease in RAD51 foci in haploids at the 2-cell stage compared to diploids, despite the increase in _γ_H2AX foci (**Fig. 4H**).

### Nuclear-cytoplasmic ratio, not ploidy, determines cytokinesis in mouse embryos

In considering possible differences between haploid and diploid ESCs that could contribute to diploidization, the most striking is the sheer difference in oocyte size relative to nuclear material. The ratio of nuclear material to cytoplasmic content, commonly referred to as the N-C ratio, differs, with haploids containing more cytoplasm. In cleavage development post-fertilization, cell volumes are halved with each successful division. As the size of an embryo’s cells decreases with each successful division during early development, we decided to compare N-C ratio relative to the physiologically normal ratio at the 2-cell and 4-cell stages.

Determining whether the instability of the haploid state could be altered by changes to the N-C ratio was undertaken through several key manipulations. Diploid mouse embryos were made haploid by the enucleation of a single nucleus in late G2 phase, just before the first cell division (labeled 2n->1n). These embryos then continued to develop as haploids until analysis at the second mitosis (**Fig. 5A**). The reduced N-C ratio is representative of a haploid cell, but the actual state of haploidy is confined to the second cell cycle instead of both first and second cell cycles, unlike originally haploid embryos. This addresses the question whether haploidy in the first cell cycle amplifies the abnormalities in the second cell cycle. Conversely, haploids were subjected to electrofusion at the 2-cell stage to produce diploids (labeled 1n->2n) (**Fig. 5A**). These newly formed diploids possess the N-C ratio of a haploid cell, as the electrofusion fuses the cytoplasm of both cells, resulting in a larger cell than typical at this stage. Additionally, 2n->1n haploids were also subjected to electrofusion to return those cells to their original diploid state (labeled 2n->1n->2n), such that they only existed as haploids during the first mitosis, but not before or thereafter (**Fig. 5A**). At the first mitosis, 2n->1n embryos showed no increase in chromosome segregation errors relative to controls. At the second mitosis in 2n->1n, 1n->2n, and 2n->1n->2n embryos all exhibited a high degree of anaphase abnormalities, comparable to what was observed in the unmanipulated haploid group (**Fig. 5B, C**). 2n->1n and 1n->2n embryos were also subjected to midbody analysis. At the second mitosis, unmanipulated haploids, as well as 1n->2n and 2n->1n embryos exhibited a dramatic decrease in the formation of a midbody, indistinguishable to unmanipulated haploids (**Fig. 5C, D**). Being diploid but with the decreased N-C ratio of a haploid, resulted in these cells also having anaphase abnormalities and frequent failure of midbody formation.

**Figure 5.**
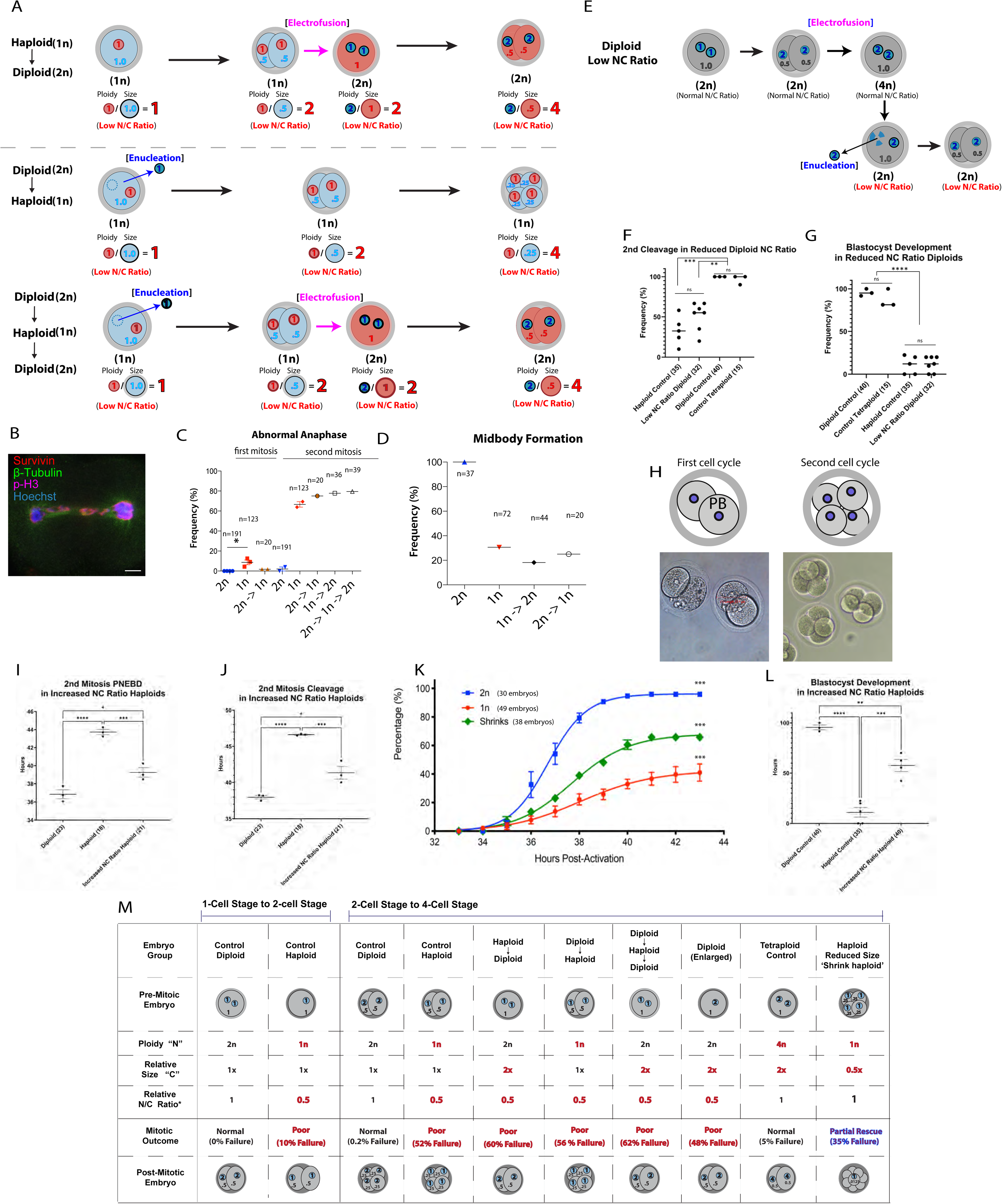
Diploids with the N-C ratio of a haploid undergo abnormal cytokinesis and fail development. **A**) Schematic illustrating manipulations performed to alter N-C ratio in haploid and diploid mouse embryos. **B**) Immunostaining illustrating abnormal anaphase in 1n->2n manipulated embryo. **C**) Frequency of chromosome segregation errors identified by anaphase abnormalities in the first and second mitoses. **D**) Frequency of midbody formation quantified by Survivin and β- tubulin immunostaining in manipulated and unmanipulated embryos in the second mitosis. **E**) Schematic illustrating manipulations performed to alter N-C ratio of diploid and tetraploid states in mouse embryos. **F**) Quantification of successful cleavages observed at the second mitosis in control diploid, control tetraploid, haploid, and “enlarged” diploid mouse embryos. **G**) Quantification of blastocyst development in indicated groups. **H**) Representative images of haploid and diploid embryos of low and normal N-C ratios. The second polar body is of equal size as the main egg, resulting in a 2-cell like embryo at the 1-cell stage. (**I**) Quantification of pronuclear envelope breakdown timing at the second mitosis determined by live-cell imaging data taken in 10-minute intervals. Each point corresponds to a single cell. (**J**) Quantification of cleavage timing at the second mitosis by live-cell imaging data taken in 10-minute intervals. Each point corresponds to a single cell. **K**) Cleavage timing and percentage of cleavage in haploids and shrinks as well as controls. **L**) Quantification of successful blastocyst development. Each point corresponding to an embryo. I-L: statistics using One-way ANOVA. **M**) Summary diagram of the different manipulations. N-C ratio of 1 is the physiologically normal ratio for diploid cells at the respective developmental stage.

Building upon these findings, additional manipulations were performed in which diploids were fused at the 2-cell stage to produce tetraploid cells, which would have an unchanged N-C ratio given both the ploidy and cell size of these cells increases proportionally. This also controls for any effects of cell fusion. In some fused tetraploid embryos, one of the two nuclei was removed, resulting in diploids with larger, fused cell size and thus a low N-C ratio (**Fig. 5E**). Diploid embryos with a low N-C ratio performed identically to haploid embryo groups with a low N-C ratio, whereas the control tetraploid embryos performed similarly to control diploid embryos.

Embryos with a low N-C ratio, whether haploid or diploid, exhibited significantly poorer cleavage rates at the second mitosis compared to embryos with a normal N-C ratio, with no statistical differences between diploid and tetraploid embryos of a normal N-C ratio (**Fig. 5F**). Furthermore, diploids with a low N-C ratios exhibited poor development indistinguishable from haploids (**Fig. 5G**).

### Increasing Nuclear-Cytoplasmic Ratio Reduces Cleavage failure and Improves Development in Haploid Mouse Embryos

To generate haploid mouse embryos with an increased N-C ratio, symmetric extrusion of the polar body was introduced through exposure of oocytes to cytochalasin B during early stages of oocyte activation for 3h. Polar body extrusion is normally a highly asymmetric cell division; the temporary application of cytochalasin B during mitotic exit, without suppression of polar body extrusion can render this division symmetric. The symmetric division leads to haploid embryos with cell sizes half of the typical 2-cell stage haploid and diploid mouse embryos. As such, these cytoplasm-reduced embryos are referred to here as 1n normal N-C ratio embryos. At the 1-cell stage, these appear as 2-cell embryos, and at the 2-cell stage, 4 cells are apparent (**Fig. 5H**). This manipulation yields embryos that possess a smaller cell size and thus higher N-C ratio compared to unmanipulated haploid embryos and thus a higher N-C ratio. We termed these ‘shrink haploids’. Pronuclear envelope breakdown during the second mitosis occurred earlier in ‘shrink haploids’ with normal N-C ratio embryos compared with unmanipulated haploid embryos (4h) (**Fig. 5I**). The time until the second mitotic cleavage was also advanced, (**Fig. 5J**), and overall cleavage was improved (**Fig. 5K**). Furthermore, blastocyst development was improved (50%) compared to haploid embryos (25%) (**Fig. 5L**). These data show that the proportions in cell volume versus nuclear content is an important determinant of mitotic cleavage, cytokinesis, genome stability and developmental potential.

## DISCUSSION

Cells of different ploidy contain the same genome, but dramatically different developmental potential. Haploid embryos fail to develop, though at the cellular level, they can activate all required developmental programs, giving rise even to haploid neurons (Sagi et al., 2016). And tetraploid cells are developmentally biased towards extraembryonic tissues. The cellular and molecular mechanisms for how differences in ploidy result in different developmental outcomes are not well understood.

The work presented here shows that haploidy results in abnormal mitosis characterized by segregation errors, and a failure to form a midbody, followed by failure of abscission, binucleation and diploidization, as well as elevated DNA damage, and reduced developmental potential. These abnormalities are at least in part driven by an abnormal ratio between genome content and cell size, since reducing this ratio in diploids reproduces these defects. In haploids with a corrected ratio of nucleus to cytoplasm, phenotypic rescue is observed, though it is not complete. This suggests stage specific requirements of ploidy relative to plasma membrane area to allow normal spindle assembly and cytokinesis.

A key component of successful cytokinesis is the appropriate formation and localization of the midbody, which we visualized through antibody staining for Survivin and AuroraB, components of the chromosome passenger complex (CPC). The CPC regulates bipolar attachment to the spindle, chromosome segregation, cytokinesis and cytoplasmic segregation, preventing a ploidy increase (van der Waal et al., 2012). In mouse haploid embryos, components of the CPC fail to relocalize to a midbody and abscission fails, providing a mechanism for diploidization. Another study in human embryos showed that RhoA, a small GTPase involved in cytokinesis, fails to localize to the midzone in haploids, leading to abscission failure (Leng et al., 2017). Surprisingly however, even when there is accurate bipolar segregation of sister chromatids, formation of a midbody can fail. The upstream mechanisms leading to defects in metaphase alignment, mislocalization of CPC components, and the formation of a midbody in haploids are not well understood.

One of the possible causes of spindle problems are DNA damage in the preceding cell cycle. Haploid cells also show regions of delayed DNA synthesis (Edwards et al., 2021), which might affect chromosome segregation. However, DNA damage occurs primarily after the 2-cell stage, and after major problems are observed in the second mitosis second mitosis. Furthermore, the chromosome segregation errors in haploid mouse embryos at primarily whole chromosome errors, suggesting that stalled forks and their breakage at mitotic entry are not a primary cause of these abnormalities. Nevertheless, increased DNA damage and lower genome stability is one of the effects of haploidy. Interestingly, an enlargement of somatic diploid cells also results in increased DNA damage (Manohar et al., 2023). When somatic cells are enlarged by extending G1 phase, the recruitment of 51BP1 to DSBs is impaired. In this study, we found a reduction of Rad51 foci, despite an overall increase in DNA damage. There are other parallel phenotypes between enlarged somatic cells and haploid embryos and enlarged embryos. Enlarged somatic cells enter mitosis with a delay, spend more time in mitosis, and often fail to undergo cytokinesis, resulting in binucleate cells (Manohar et al., 2023). A mechanistic understanding of how cell enlargement results in mitotic abnormalities is incomplete.

Several studies have reported advances for increasing haploidy in culture. These include stimulating premature entry into mitosis by Wee1 inhibition (Takahashi et al., 2014). Wee1 kinase was originally identified in yeast when loss of function mutations caused yeast to enter mitosis at a smaller size. This again points to cell size as an important factor in determining the stability of the haploid state. However, another study reported that CKD1 inhibition also helped to increase haploidy (He et al., 2017). This manipulation would be expected to increase cell size by lengthening the cell cycle, though this was not directly examined. Rock inhibition, which inhibits cell death of dissociated pluripotent stem cells after passaging, also helped to maintain haploidy (He et al., 2017). This effect may in part be due to an overall improved cell culture rather than specific to haploidy. Another study reported that reducing oxygen tension increased haploidy (Zhang et al., 2020). However, in this study, we found no effect of reduced oxygen tension of haploid mouse embryo development or frequency of mitotic abnormalities.

Another study reported that low concentrations of microtubule stabilizing compounds such as paclitaxel selectively removes diploids and tetraploid cells and enriches for haploids (Olbrich et al., 2019). Conversely, statins, inhibitors of cholesterol synthesis, accelerated the loss of haploid cells relative to diploid cells in mixed cultures, pointing to the relevance of the plasma membrane (Olbrich et al., 2019). This is consistent with our findings that not cytoplasmic volume but cell- or plasma membrane area relative to genomic content is the primary determinant of stable ploidy.

The sizes of nucleus, cytoplasm and mitotic spindle change dramatically during preimplantation development, from the 1-cell to the blastocyst stage. Chromosome condensation also occurs in proportion to cell and nuclear size (Ladouceur et al., 2015). Thus, relevant proportions of cell size, nuclear size, spindle and chromatid length are stage- and species- specific. At the 1-cell stage, a low nucleus to cytoplasm ratio is tolerated, but the same ratio at the 2-cell stage results in mitotic failure. The mechanisms determining this stage-specific tolerance are not known. Human or cow oocytes are over twice as large as mouse oocytes, but this does not intrinsically result in abnormal cytokinesis. Interestingly, the development of haploids is only modestly compromised in human compared to normal diploids (∼25% development in haploids instead of ∼50% in diploids (Sagi et al., 2016), contrasting the findings here in mice where development drops from 95% to ∼10-20%. Diploidization occurs primarily after embryonic genome activation, as shown here at the division from 2 to 4-cells in haploid mouse embryos. Notably in human, zygotic genome activation occurs in the third and fourth cell cycle, potentially underlying the greater tolerance to haploidy in human than in mice.

The data presented here provide novel aspects of the biology as to why we exist as diploid organisms. While human and other mammalian stem cells can exist in a haploid state, they do not give rise to a complex organism. Genome instability in haploids points to the delicate requirements for progression through the cell cycle, and adequate ratios of cytoplasm and nucleus or mitotic spindle. Haploid stem cells and haploid mouse embryos provide a model system to study the mechanisms determining stable ploidy. The work presented here will enable future studies on the molecular interactions between genome content, cell size, plasma membrane area, chromosome condensation, spindle size and function, DNA replication and DNA damage, and the specific signals resulting in ploidy change with implications spanning development, evolution, and disease.

## Acknowledgements

This work was supported by NYSTEM grants #C32564GG, the US Israel Binational Science Foundation grant # 2015089. We thank Julie Canman, Alberto Ciccia, Rodney Rothstein, and Hynek Wichterle for critical reading and input in experimental design and data interpretation, Robin Goland for help with human subjects research, and Rudolph L. Leibel for project input in institutional presentations.

## Author contribution

Project design and lead was performed by MVZ, conducting all studies, including cell culture, live cell imaging, flow cytometry, embryo studies, with contributions from others as below.

Oocyte donor consent and human embryo and gamete collection and preparation was performed by Dr. RL and the CUFC embryology team. RP provided contribution to human embryo studies. Immunofluorescent staining and confocal microscopy of human embryos was performed by LCG. BWGA was performed by SA and analysis by SX. ADLA contributed to genome amplifications and MD to analysis. DE provided embryology, micromanipulation, supervision and assistance to experimentation, data interpretation and manuscript writing.

## METHODS

### Mouse embryo activation

Activation of haploid and diploid mouse embryos was induced using ionomycin as a calcium ionophore for primary activation, followed by continued activation through puromycin-induced inhibition of protein synthesis. Diploid embryos were generated using a 5-hour exposure to cytochalasin B to prevent actin polymerization and thus asymmetric polar body extrusion, resulting in a single cell embryo with a diploid genomic content.

### Derivation of control diploid embryonic stem cell lines using defined media

Before characterizing and comparing haploid and isogenic human embryonic stem cells, additional diploid cell lines were derived with *in vitro* fertilization as is typically done. 21 blastocysts of variable quality, which were all consented for use in scientific research and were not designated for use in human reproduction. Excess frozen human cleavage stage embryos not designated for reproduction were thawed and incubated in Quinn’s Advantage Thaw Kit under standard culture procedures. Cleavage stage embryos and day 5 blastocysts were cultured until day 6 or the expanded blastocyst stage. The inner cell mass of day 6 blastocysts was plated and expanded (**Fig. S1A**). G-banding karyotype analysis was performed, and karyotypically normal stem cells were further characterized to ensure normal morphology and pluripotency (**Fig. S1B**). Morphology appeared normal, yielding tightly bordered colonies of embryonic stem cells throughout 20 passages in culture (**Fig. S1C**). Immunofluorescent staining was performed for pluripotency markers TRA-1-60 and SSEA-4 (**Fig. S1D**). Pluripotency was further confirmed with flow cytometry for double-positive cells for TRA-1-60 and SSEA-4 (**Fig. S1E**). Cells were also made to form embryoid bodies, which were immunostained for markers of the three germ layers, endoderm, mesoderm, and ectoderm (**Fig. S1F**). These data indicate that the IVF-derived lines were indeed embryonic stem cell lines and were incorporated in subsequent analyses as diploid controls.

### Immunostaining

Primary antibody incubation was performed on cells fixed in 4% PFA in PBS (Santa Cruz Biotechnology, sc-281692) for 15 minutes at room temperature, permeabilized in 0.1% Triton® X-100 (Sigma Aldrich, T8787) in PBS for 30 minutes at room temperature, and blocked in 10% FBS (Gemini Bio-Products, 900-108) in PBS for 30 minutes at room temperature. Incubation was performed at the indicated dilution in 10% FBS in PBS for 4 hours at room temperature.

Primary antibodies used include beta-tubulin antibody (clone AA2, Millipore 05-661, dilution 1:1,000), γH2AX (Cell Signaling 2257S)1:1,000, Phospho-Histone H3 (Thermo Fisher 06-570) 1:,1000, AuroraB 1:500 (Abcam an2254), 53BP1 (BD-612522) 1:1,000), Rad51 (EDM Millipore PC130) 1:200, Centromere (Antibodies Inc.) 1:50, RPA32 (Cell Signaling Technologies 2208S) 1:200, and Survivin (Cell Signaling Technologies 71G4B7) 1:200.

After 3 washes with 10% FBS in PBS, secondary antibody incubation was performed for 1 hour at room temperature, diluted in 10% FBS in PBS. Secondary antibodies include Invitrogen 488, 555, and 647 anti mouse (A21202), (A31570), and (A-31571) at 1:500, Invitrogen 488, 555, and 647 anti mouse (A32731), (A-31572), and (A32733) at 1:500, Invitrogen 488, 555, and 647 anti rat (A48262), (A48270), and (A- A32733) at 1:500, and Hoechst 33342 was used for DNA staining at 5 μg/mL (Life Technologies H3570).

### Cell cycle analysis & flow cytometry

For cell cycle analysis including S-phase, cells were incubated in pre-warmed media with 10 μM EdU (Life Technologies, C10337) for 30 minutes followed by a pre-warmed media change without EdU. Negative control cells were not exposed to EdU. Cells were incubated until the desired time point, upon which media was aspirated, cells were rinsed with PBS (LifeTechnologies 14190-250), trypsinized with TrypLE Express (LifeTechnologies, 12605036) for 5 minutes, and suspended in 4% PFA in PBS (Santa Cruz Biotechnology, sc-281692) for 15 minutes. Cells were pelleted and suspended in 0.1% Triton® X-100 (Sigma Aldrich, T8787) for 30 minutes. Cells were pelleted and suspended in 10% FBS (Gemini Bio-Products, 900-108) in PBS for 30 minutes, followed by optional immunostaining methods (described above) for antibody inclusion. Prior to analysis, cells were incubated in 10 μg/mL Hoechst 33342 (Life Technologies, H3570) for 30 minutes and washed once with 10% FBS in PBS.

For live-cell analysis, cells were suspended and pelleted as described above, and suspended in medium with 10 μM Flow cytometry was performed using the ZE5 Biorad machine or the FACS-Aria machine at the Columbia University Stem Cell Initiative flow cytometry core. Populations were gated first for cells, followed by gating for single cells. Cells were suspended in media containing 10% FBS in PBS throughout the analysis. Live cells were kept on ice during transportation and analysis.

Analysis was performed using FloJo software (BD Biosciences).

### Mouse Strains, Oocytes

5–7-week-old B6D2F1/J female mice were used for obtaining mouse oocytes (Jackson Laboratories stock 10006). Mice were injected with 5 IU/mouse of pregnant mare serum gonadotropin (ProSpec, HOR-272), followed by an injection of 5 IU/mouse of human chorionic gonadotropin (EDM Millipore, 230734) within 48hr later. 14-16hr post hCG injection, mice were euthanized and oviducts were collected. Oocytes were removed from oviducts and equilibrated in Global Total (LifeGlobal, H5GT-030). Activation was performed by 5min exposure to ionomycin (Sigma-Aldrich, I3909) in Global Total, followed by incubation in Global Total with puromycin (Life Technologies, A11138-03) for haploids or puromycin and cytochalasinB (Sigma Aldrich, C2743) for diploids. Manipulations were performed on an inverted Olympus IX73 equipped with a Narishige micromanipulator. Images were taken using an Olympus IX73 equipped with an Olympus DP80 camera or a Zeiss 710 confocal microscope at indicated time points.

All animal research was reviewed and approved by the Columbia University IACUC, and performed in accordance with animal use guidelines and applicable ethical regulations.

### Human gamete donation, embryos

Fresh oocytes were obtained from women ages 22-29 years who consented to undergo an oocyte donor cycle to donate oocytes to stem cell research. Metaphase II (MII) oocytes were manipulated within 4 hours of donation, while immature oocytes were cultured in global total media for up to 48 hours and then used if they became a MII oocyte. MII oocytes were either activated by an electrical pulse and puromycin to create parthenogenetic (P) embryos, or enucleated and injected with sperm via intracytoplasmic sperm injection to create androgenetic (A) embryos. Embryos were incubated and the morphology was assessed daily. Embryos with poor morphology or delayed growth were fixed in 4% paraformaldehyde, stained for human centromere protein A and analyzed by confocal microscopy to count centromeres for ploidy determination. Embryos that became a fair or good quality blastocyst by Day 6 underwent trophectoderm (TE) biopsy that was used for ploidy evaluation, while the inner cell mass was plated to grow a stem cell line.

### Stem cell derivation

Stem cells were derived after trophectoderm biopsy and plating for the inner cell mass as previously described (Yamada et al., 2014). Briefly, mural trophectoderm was ablated using laser-assisted pulses 400 μs, 100% intensity (Hamilton Thorne, Chen et al., 2009). This method spares polar trophectoderm, which usually results in trophectoderm growth which is then ablated with additional pulses. Derivation of uniparental lines was performed on irradiated MEFs (GlobalStem) in KO-DMEM with 25% KO-SR with 10 μM Rock inhibitor Y-27632, 10ng/mL bFGF and 2% ES grade FBS. Outgrowths were allowed to grow for two weeks until manual passaging and FBS is gradually phased in partial media changes every second or third day. For IVF ESC lines, derivation occurred on LN511E8 matrix (Takara T304) and StemFit medium (Ajinomoto SM-Basic-02).

### PCR for SNP identification between in-house derived IVF lines

PCR was performed with the following primer sets in the 5′ -3′ orientation: CCTGAGCTGGAGCCTGGGAGCC … GGCGGGGAGAAGGGCTCTTAG CCTGAAACCATTTTGGCTAGA … CTATTACTGGTAGAACTGATG ATATTAATATTTGCTAATAAT … TTATGGATATGTTAATTTGCT ATGTGAAGCAGGTCCAAATAT … AATTTACAATCACTCAATCTT

### H2B-GFP haploid ESCs

Plasmid CD615B-1-SBI Lentiviral vector was from SBI System Biosciences and H2B-GFP was cloned using competent cells (Invitrogen #18258012). Lentiviral particles were transfected into HEK293 cells and collected after 24 hours. Stem cells were exposed to virus for 6 hours and cultured for 72 hours post-exposure. Selection was performed by flow cytometry to select for cells expressing GFP.

### Stem cell culture

Passaged stem cells a cultured with StemFlex media on Geltrex. Upon reaching 70% confluency, cultures were passaged at a ratio of 1:10, or cryopreserved in a solution of freezing media containing 40% FBS (Company) and 10% DMSO (Sigma Aldrich). Passaging was performed by TrypLE dissociation to small clusters of cells, and plated in media containing 10 μM Rock inhibitor Y-27632 was added to media and removed within 24-48 h. For later passage cells (> passage 10), Rock inhibitor was omitted. All embryo and ESC research was reviewed and approved by the Columbia University Embryonic Stem Cell Committee and the Institutional Review Board.

### Small molecule inhibitors

CHK1/2 (Selleckchem, AZD7762), CHK1 (Tocris, LY2603618). Concentrations used were determined by testing a series of concentrations and selecting the highest tolerable dose that did not impact viability of cell cultures.

### BWGA and bioinformatics analysis

Barcoded whole genome amplification (BWGA) was performed with 10X Single Cell Lysis and Fragmentation buffer (Sigma-Aldrich L1043), Klenow DNA Polymerase (NEB M0210L, New England Biolabs), AMPure XP beads (Beckman Coulter A63880), NEBNext Ultra II DNA Library Prep Kit for Illumina (New England Biolabs). Universal primers and barcodes were provided by Timour Baslan of Memorial Sloan Kettering. Analysis was performed using R software. Sequencing data were aligned to the human genome (hg19) using the Burrows-Wheeler Aligner^1^. The aligned genome was partitioned into 100kb bins for copy number calling. The plots and segment files were generated using the QDNAseq package^2^.

For randomly generated control comparisons, intervals within mm10 genome assembly were randomly generated using bedtools v2.30.01^3^. The plots and segment files were generated using the QDNAseq package2. The mm10 genome assembly chromosome length file was downloaded from: http://hgdownload.cse.ucsc.edu/goldenpath/mm10/bigZips/mm10.chrom.sizes

### BWGA Methods References

1. Li H. (2013) Aligning sequence reads, clone sequences and assembly contigs with BWA-MEM. arXiv:1303.3997v2 [q-bio.GN].
2. Scheinin I, Sie D, Bengtsson H, van de Wiel MA, Olshen AB, van Thuijl HF, van Essen HF, Eijk PP, Rustenburg F, Meijer GA, Reijneveld JC, Wesseling P, Pinkel D, Albertson DG, Ylstra B (2014). “DNA copy number analysis of fresh and formalin-fixed specimens by shallow whole-genome sequencing with identification and exclusion of problematic regions in the genome assembly.” Genome Research, 24, 2022–2032.
3. Quinlan AR, Hall IM, 2010. BEDTools: a flexible suite of utilities for comparing genomic features. Bioinformatics. 26, 6, pp. 841–842.

## Data Availability

Single cell sequencing data are available at SRA under accession number PRJNA874697 and PRJNA1292278 using reviewer link https://dataview.ncbi.nlm.nih.gov/object/PRJNA1292278?reviewer=i35hckomi2g0r4s3a6o2o5q7e4

---

**Figure S1.**
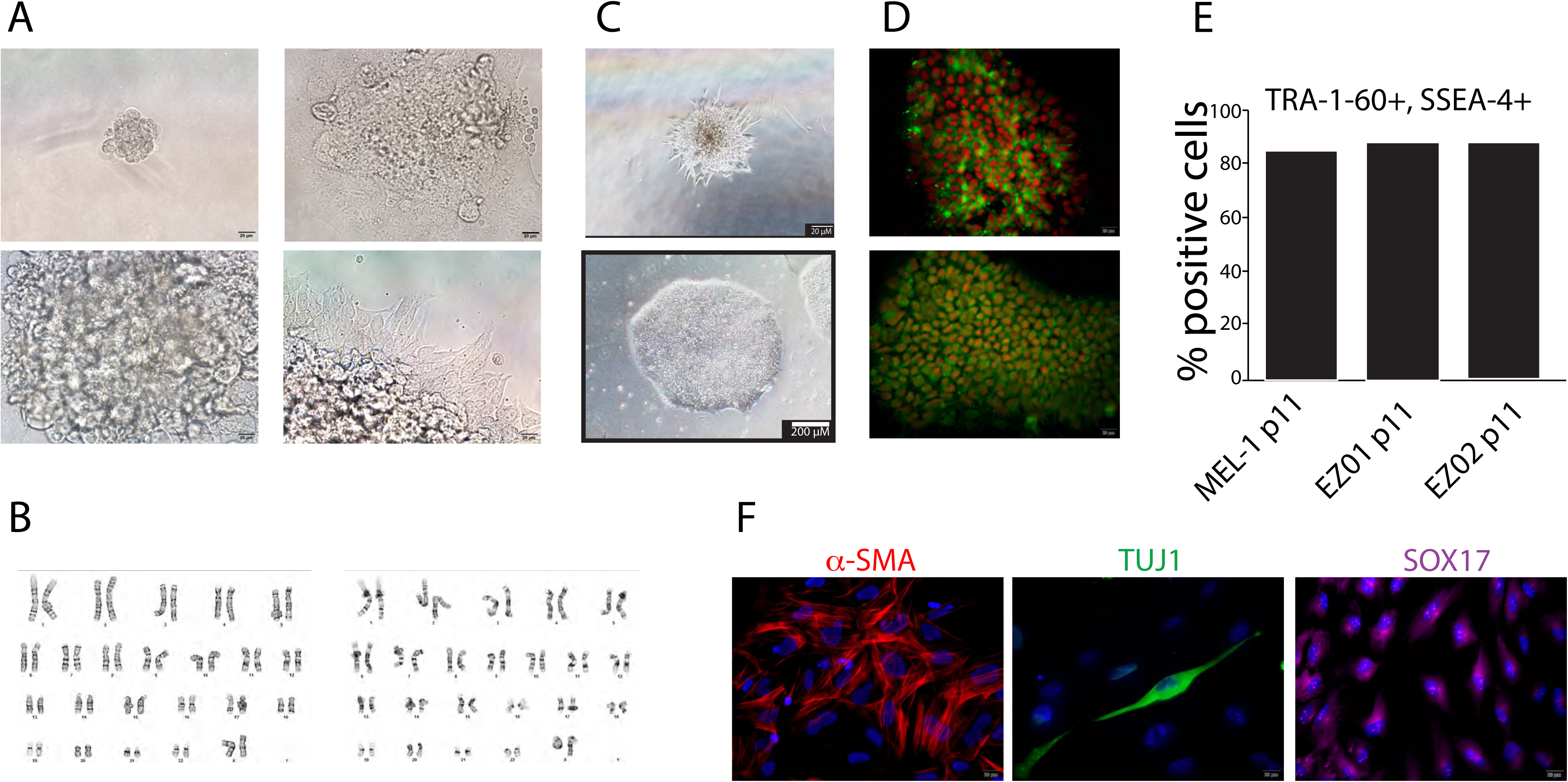
Derivation of new IVF-derived diploid human embryonic stem cell lines under defined conditions. (**A**). Overview of derivation process across multiple days. ESCs migrating from inner cell mass outgrowth indicated by black arrow. Scale bar = 20 _μ_m. (**B**) Karyotype analysis of cell lines EZ01 and EZ02 indicating normal chromosome content. (**C**) Normal morphology observed under brightfield microscopy at passage 2 (top) and passage 20 (bottom) for ESC line EZ01. Scale bar = 200 _μ_m. (**D**) Immunofluorescent staining of pluripotency markers. Scale bar = 50 _μ_m. (**E**) Quantification of flow cytometry data analyzing double-positive population frequency for TRA-1-60 and SSEA-4 in derived lines and Mel-1 ESCs as a comparison. (**F**) Immunofluorescent staining of three germ layer markers SMA, Tuj1, and SOX17. Scale bar = 20 _μ_m.

**Table S1.** | Chromosomal content of nuclei and blastomeres of haploid embryos

## References

1. Celton-Morizur, S., Merlen, G., Couton, D., Margall-Ducos, G. & Desdouets, C. 2009. The insulin/Akt pathway controls a specific cell division program that leads to generation of binucleated tetraploid liver cells in rodents. J Clin Invest, 119, 1880–7.

2. Davoli, T. & De Lange, T. 2011. The causes and consequences of polyploidy in normal development and cancer. Annu Rev Cell Dev Biol, 27, 585–610.

3. Ding, C., Huang, S., Qi, Q., Fu, R., Zhu, W., Cai, B., Hong, P., Liu, Z., Gu, T., Zeng, Y., Wang, J., Xu, Y., Zhao, X., Zhou, Q. & Zhou, C. 2015. Derivation of a Homozygous Human Androgenetic Embryonic Stem Cell Line. Stem Cells Dev, 24, 2307–16.

4. Edwards, M. M., Zuccaro, M. V., Sagi, I., Ding, Q., Vershkov, D., Benvenisty, N., Egli, D. & Koren, A. 2021. Delayed DNA replication in haploid human embryonic stem cells. Genome Res.

5. Elling, U., Taubenschmid, J., Wirnsberger, G., O’malley, R., Demers, S. P., Vanhaelen, Q., Shukalyuk, A. I., Schmauss, G., Schramek, D., Schnuetgen, F., Von Melchner, H., Ecker, J. R., Stanford, W. L., Zuber, J., Stark, A. & Penninger, J. M. 2011. Forward and reverse genetics through derivation of haploid mouse embryonic stem cells. Cell Stem Cell, 9, 563–74.

6. Freimann, R. & Wutz, A. 2017. A fast and efficient size separation method for haploid embryonic stem cells. Biomicrofluidics, 11, 054117.

7. Guidotti, J. E., Brégerie, O., Robert, A., Debey, P., Brechot, C. & Desdouets, C. 2003. Liver cell polyploidization: a pivotal role for binuclear hepatocytes. J Biol Chem, 278, 19095–101.

8. Guo, A., Huang, S., Yu, J., Wang, H., Li, H., Pei, G. & Shen, L. 2017. Single-Cell Dynamic Analysis of Mitosis in Haploid Embryonic Stem Cells Shows the Prolonged Metaphase and Its Association with Self-diploidization. Stem Cell Reports, 8, 1124–1134.

9. He, Z. Q., Xia, B. L., Wang, Y. K., Li, J., Feng, G. H., Zhang, L. L., Li, Y. H., Wan, H. F., Li, T. D., Xu, K., Yuan, X. W., Li, Y. F., Zhang, X. X., Zhang, Y., Wang, L., Li, W. & Zhou, Q. 2017. Generation of Mouse Haploid Somatic Cells by Small Molecules for Genome-wide Genetic Screening. Cell Rep, 20, 2227–2237.

10. Hirabayashi, M., Hara, H., Goto, T., Takizawa, A., Dwinell, M. R., Yamanaka, T., Hochi, S. & Nakauchi, H. 2017. Haploid embryonic stem cell lines derived from androgenetic and parthenogenetic rat blastocysts. J Reprod Dev, 63, 611–616.

11. Kaufman, M. H., Robertson, E. J., Handyside, A. H. & Evans, M. J. 1983. Establishment of pluripotential cell lines from haploid mouse embryos. J Embryol Exp Morphol, 73, 249–61.

12. Ladouceur, A.-M., Dorn, J. F. & Maddox, P. S. 2015. Mitotic chromosome length scales in response to both cell and nuclear size. Journal of Cell Biology, 209, 645–652.

13. Leeb, M. & Wutz, A. 2011. Derivation of haploid embryonic stem cells from mouse embryos. Nature, 479, 131–4.

14. Leng, L., Ouyang, Q., Kong, X., Gong, F., Lu, C., Zhao, L., Shi, Y., Cheng, D., Hu, L., Lu, G. & Lin, G. 2017. Self-diploidization of human haploid parthenogenetic embryos through the Rho pathway regulates endomitosis and failed cytokinesis. Scientific Reports, 7, 4242.

15. Leone, M., Musa, G. & Engel, F. B. 2018. Cardiomyocyte binucleation is associated with aberrant mitotic microtubule distribution, mislocalization of RhoA and IQGAP3, as well as defective actomyosin ring anchorage and cleavage furrow ingression. Cardiovasc Res, 114, 1115–1131.

16. Li, W., Li, X., Li, T., Jiang, M. G., Wan, H., Luo, G. Z., Feng, C., Cui, X., Teng, F., Yuan, Y., Zhou, Q., Gu, Q., Shuai, L., Sha, J., Xiao, Y., Wang, L., Liu, Z., Wang, X. J., Zhao, X. Y. & Zhou, Q. 2014. Genetic modification and screening in rat using haploid embryonic stem cells. Cell Stem Cell, 14, 404–14.

17. Manohar, S., Estrada, M. E., Uliana, F., Vuina, K., Alvarez, P. M., De Bruin, R. a. M. & Neurohr, G. E. 2023. Genome homeostasis defects drive enlarged cells into senescence. Mol Cell, 83, 4032–4046.e6.

18. Olbrich, T., Vega-Sendino, M., Murga, M., De Carcer, G., Malumbres, M., Ortega, S., Ruiz, S. & Fernandez-Capetillo, O. 2019. A Chemical Screen Identifies Compounds Capable of Selecting for Haploidy in Mammalian Cells. Cell Reports, 28, 597–604.e4.

19. Qu, C., Yan, M., Yang, S., Wang, L., Yin, Q., Liu, Y., Chen, Y. & Li, J. 2018. Haploid embryonic stem cells can be enriched and maintained by simple filtration. J Biol Chem, 293, 5230–5235.

20. Sagi, I. & Benvenisty, N. 2017. Haploidy in Humans: An Evolutionary and Developmental Perspective. Dev Cell, 41, 581–589.

21. Sagi, I., Chia, G., Golan-Lev, T., Peretz, M., Weissbein, U., Sui, L., Sauer, M. V., Yanuka, O., Egli, D. & Benvenisty, N. 2016. Derivation and differentiation of haploid human embryonic stem cells. Nature, 532, 107–11.

22. Sagi, I., De Pinho, J. C., Zuccaro, M. V., Atzmon, C., Golan-Lev, T., Yanuka, O., Prosser, R., Sadowy, A., Perez, G., Cabral, T., Glaser, B., Tsang, S. H., Goland, R., Sauer, M. V., Lobo, R., Benvenisty, N. & Egli, D. 2019. Distinct Imprinting Signatures and Biased Differentiation of Human Androgenetic and Parthenogenetic Embryonic Stem Cells. Cell Stem Cell, 25, 419–432.e9.

23. Takahashi, S., Lee, J., Kohda, T., Matsuzawa, A., Kawasumi, M., Kanai-Azuma, M., Kaneko-Ishino, T. & Ishino, F. 2014. Induction of the G2/M transition stabilizes haploid embryonic stem cells. Development, 141, 3842–7.

24. Van Der Waal, M. S., Hengeveld, R. C. C., Van Der Horst, A. & Lens, S. M. A. 2012. Cell division control by the Chromosomal Passenger Complex. Experimental Cell Research, 318, 1407–1420.

25. Yang, H., Shi, L., Wang, B. A., Liang, D., Zhong, C., Liu, W., Nie, Y., Liu, J., Zhao, J., Gao, X., Li, D., Xu, G. L. & Li, J. 2012. Generation of genetically modified mice by oocyte injection of androgenetic haploid embryonic stem cells. Cell, 149, 605–17.

26. Zhang, X. M., Wu, K., Zheng, Y., Zhao, H., Gao, J., Hou, Z., Zhang, M., Liao, J., Zhang, J., Gao, Y., Li, Y., Li, L., Tang, F., Chen, Z. J. & Li, J. 2020. In vitro expansion of human sperm through nuclear transfer. Cell Res, 30, 356–359.

